# Insights into DNA substrate selection by APOBEC3G from structural, biochemical, and functional studies

**DOI:** 10.1101/235481

**Authors:** Samantha J. Ziegler, Chang Liu, Mark Landau, Olga Buzovetsky, Belete A. Desimmie, Qi Zhao, Tomoaki Sasaki, Ryan C. Burdick, Vinay K. Pathak, Karen S. Anderson, Yong Xiong

## Abstract

Human apolipoprotein B mRNA-editing enzyme-catalytic polypeptide-like 3 (A3) proteins are a family of cytidine deaminases that catalyze the conversion of cytidine to uridine in single-stranded DNA (ssDNA). A3 proteins act in the innate immune response to viral infection by mutating the viral ssDNA. One of the most well-studied human A3 family members is A3G, which is a potent inhibitor of HIV-1. Each A3 protein prefers a specific substrate sequence for catalysis - for example, A3G deaminates the third cytidine in the CC**<u>C</u>**A sequence motif. However, the interaction between A3G and ssDNA is difficult to characterize due to poor solution behavior of the full-length protein and loss of DNA affinity of the truncated protein. Here, we present a novel DNA-anchoring fusion strategy, which we have used to capture an A3G-ssDNA interaction. We characterized an A3G-DNA binding pocket that is important for the enzyme to scan the DNA for its hotspot. The results provide insights into the mechanism by which A3G selects and deaminates its preferred substrates and help define how A3 proteins are tailored to recognize specific DNA sequences. This knowledge contributes to a better understanding of the mechanism of DNA substrate selection by A3G, as well as A3G antiviral activity against HIV-1.

## Introduction

Apolipoprotein B mRNA-editing enzyme-catalytic polypeptide-like 3G (APOBEC3G or A3G) is a human host restriction factor that inhibits human immunodeficiency virus type 1 (HIV-1), murine leukemia virus, and equine infectious anemia virus (1–5) primarily through its cytidine deaminase activity on the viral minus-strand DNA (-ssDNA). A3G is one of the seven APOBEC3 (A3) proteins that inhibits replication of a diverse set of viruses (4, 6, 7). The human cytidine deaminase superfamily members have highly conserved protein sequences, tertiary structural folds, and catalytic mechanisms (4, 8). All A3 proteins and the closely related activation-induced cytidine deaminase (AID) contain a single catalytically active cytidine deaminase (CDA) domain. However, some family members, such as A3B, A3D, A3F, and A3G, have a second pseudocatalytic domain that retains the same tertiary fold, but are not catalytically active. A3G inhibits viral replication by preferentially deaminating cytidine residues to uridine in the viral -ssDNA during reverse transcription (7, 9–11). Specific “hotspot” sequences of DNA are targeted for deamination, with the highest preference for the 5’-CC**<u>C</u>**A sequence (where the deaminated cytidine is underlined) in the case of A3G (2, 12). Deamination of the -ssDNA by A3G leads to extensive G-to-A hypermutation in the viral genome, ultimately eliminating viral infectivity (1, 5, 12). Some studies suggest that mutations induced by A3G are occasionally sub-lethal, facilitating viral evolution and allowing the virus to develop drug resistance (3, 13). However, a recent study concluded that hypermutation is almost always a lethal event and makes little or no contribution to viral genetic variation (14). This underlines the importance of understanding the mechanism by which A3G induces mutations in the HIV-1 genome (3, 15). While the A3s’ deaminase activity is beneficial during viral restriction, this function has also been reported to promote multiple types of cancer (16). For example, the deaminase activity of A3B has been linked to mutations found in breast cancer (17), and overactive AID is believed to be responsible for non-Hodgkin’s lymphoma (18). In this regard, APOBECs are the second most probable contributors to cancer development, and thus, it is critical to uncover the mechanisms underlining the interactions between the A3 proteins and their DNA substrates (19).

Biochemical studies (20–23), as well as the apo structure of the C-terminal catalytic domain of A3G (A3G_CTD_) (20, 21, 23–25), have established a framework for A3G hotspot selectivity. Extensive biochemical and mutagenesis studies have shown that a single loop, loop 7, of A3G is responsible for selecting the base at the −1 position (the nucleotide at the 5’ side of the deamination site) of the DNA substrate hotspot (26–29). Swapping this loop into the complementary site in AID altered AID hotspot preference to that of A3G (26, 28). Although the crystal structures of A3A, A3B, and AID in complex with DNA have recently been published (30–32), a structure of a catalytically active A3G bound to its substrate has yet to be determined. The lack of structural insight into the A3G-DNA complex has limited our understanding of the molecular determinants that govern A3G specificity and selectivity for its substrate. Structural studies have been hampered by the inherent difficulty of purifying full-length A3G with sufficient solubility. The A3G N-terminus (A3G_NTD_) also readily forms soluble aggregates partly due to its propensity to bind RNA and DNA (33). Although the isolated A3G_CTD_ is able to bind and deaminate ssDNA, the loss of the A3G_NTD_ results in a drastic decrease in DNA binding affinity (21).

In this study, we used a novel protein fusion strategy to obtain structural insight into the A3G-ssDNA interaction. We captured the scanning state of the −1 nucleotide (the nucleotide at the 5’ side of deaminated cytidine) binding pocket. Our results reveal key interactions that govern substrate sequence specificity of A3G and related cytidine deaminases. Furthermore, our results reveal how these key interactions perpetuate changes throughout the A3G hotspot binding pocket. Together, this study provides a deeper understanding of A3G anti-viral activity and expands our knowledge of the deamination mechanism of related cytidine deaminases.

## Results

### Novel fusion design captures A3G_CTD_ bound to a nucleotide

Although A3G_ctd_ overcomes the solubility problem of the full-length protein and retains activity, DNA binding is weak and does not form a stable DNA-protein complex required for structural analysis. To address the problem of weak DNA binding, we designed a novel A3G_CTD_ fusion to the protein Protection of telomeres 1 (Pot1) that serves as an anchor for the DNA (experimental schematic shown in Fig 1A). Pot1 is a DNA binding protein that binds its specific cognate sequence at 1 nM affinity (34). We designed a ssDNA substrate that contains both the Pot1 cognate sequence and the A3G hotspot sequence separated by a 14-nucleotide linker sequence (Fig 1A). We confirmed that the Pot1A3G fusion binds to ssDNA using size exclusion chromatography (Fig 1B) and that the fusion construct retains its native activity towards the A3G DNA substrate (Fig 1C). These results show that Pot1 fusion may serve as a general strategy to stably anchor otherwise weakly bound DNA substrates to proteins.

**Fig 1.**
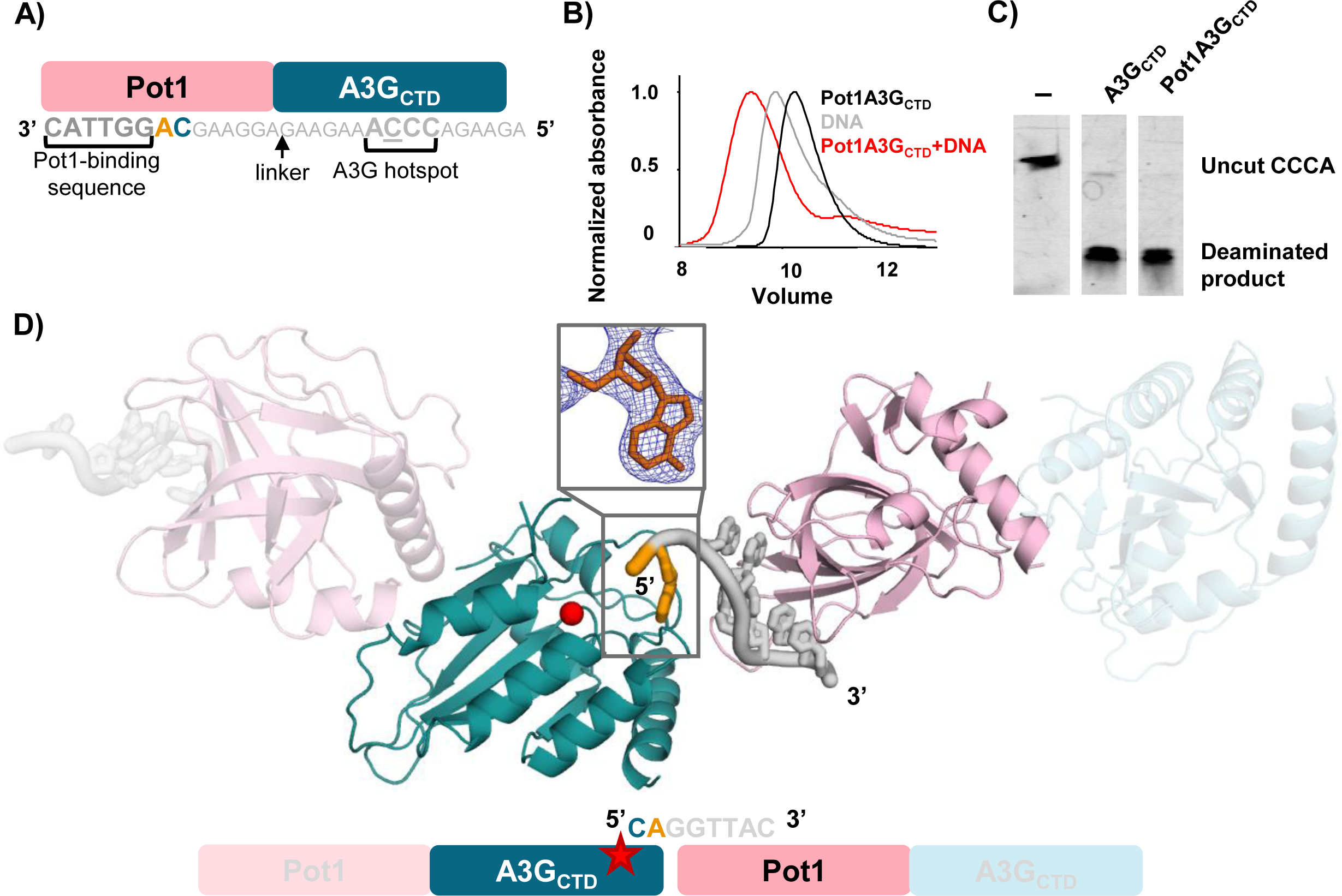
Structure of Pot1-A3G_CTD_ with ssDNA. A) Schematic of the Pot1-A3G_CTD_ fusion protein design. Pot1 (pink) is fused directly to the N-terminus of A3G_CTD_ (blue). The ssDNA contains both Pot1 and A3G binding sites: the Pot1 site in dark gray and the A3G hotspot in light gray with the linker sequence in smaller font. The resolved adenine in the −1 pocket is colored orange and the expected deaminated cytidine is blue. B) Size exclusion binding test shows that PoUA3Gctd binds to the ssDNA substrate. Pot1A3G_CTD_ alone is in black, the ssDNA is in gray, and the mixture of the two is in red. C) Deamination activity using a UDG-dependent cleavage assay. The Pot1-A3G_CTD_ fusion protein has the same deamination activity as that of A3G_CTD_. D) Schematic and structure of the Pot1-A3G_CTD_ in complex with DNA as observed in the crystal. The adenine nucleotide bound to the −1 pocket is shown in orange. Two copies of the complex observed in the asymmetric unit are shown in blue (A3G), pink (Pot1), and grey/orange (DNA). The red star (schematic) and red sphere (structure) represent the zinc ion found in the catalytic site. The inset shows the 2Fo-Fc density (1σ contour level) observed for the adenine in the −1 nucleotide-binding pocket.

Using the Pot1A3G_CTD_ fusion construct, we obtained a crystal structure of A3G_CTD_ bound to an adenine nucleotide at 2.9 Å resolution (Fig 1D and Table 1). The structure was determined in a P4_3_ space group with four independent copies of the protein in the asymmetric unit of the crystal. Both A3G_CTD_ and Pot1, together with the Pot1 cognate DNA, were clearly visible in the electron density. In the crystal, the ssDNA stretches from a Pot1 protein of one fusion complex to the A3G of the adjacent fusion complex. The rest of the ssDNA was disordered with clear electron density for only one nucleotide, adenine, located next to the catalytic site of A3G_CTD_. This fusion design allowed us to capture the structure of A3G_CTD_ bound to an adenine nucleotide in the −1 DNA binding pocket (Fig 1D inset).

**Table 1.** Data collection and refinement statistics

|  |  |
| --- | --- |
| Space group | $P 4_3$ |
| **Cell dimensions** |  |
| a, b, c (Å) | 79.07, 79.07, 266.41 |
| $\alpha$, $\beta$, $\gamma$ (°) | 90.00, 90.00, 90.00 |
| Resolution (Å) | 44-2.9 (3.0-2.9)* |
| No. reflections | 40118 |
| **Redundancy** | 3.3 (3.3) |
| $R_{\text{merge}}$ | 0.105 (0.856) |
| $R_{\text{pim}}$ | 0.064 (0.539) |
| Mean I / $\sigma$ I | 5.1 (1.1) |
| Completeness (%) | 97.5 |
| CC<sub>1/2</sub> last shell | 0.70 |
| **Refinement** |  |
| Resolution (Å) | 44-2.9 |
| No. Reflections | 32909 |
| $R_{\text{work}}$ / $R_{\text{free}}$ | 0.237/0.287 (0.30/0.37) |
| No. atoms | 11958 |
| B-factors | 88.2 |
| **RMSD in** |  |
| Bond lengths (Å) | 0.019 |
| Bond angles (°) | 1.9 |
| **Ramachandran** |  |
| Preferred | 94.6 |
| Allowed | 5.0 |
| Outliers | 0.4 |

\* Values in parentheses are for the highest-resolution shell

### A3G-adenine structure captures a scanning state of the enzyme

Our crystal structure of the Pot1-A3G_CTD_ fusion potentially captured an intermediate state of A3G during its search for a ssDNA hot spot sequence. A3G processively scans ssDNA (35) and prefers to deaminate cytidine in the context of the 5’-CC**<u>C</u>**A sequence, where the underlined C is deaminated. In this sequence, the cytidine preceding the deaminated C is referred to as the - 1 nucleotide and the adenine following the deaminated C is the +1 nucleotide (2, 12). Although we intended to crystallize the A3G_CTD_ bound to its entire hotspot region as described in figure 1a, we instead captured the A3G_CTD_ bound to an adenine in the −1 pocket (Fig 1D). Although the preferred nucleotide for A3G at the −1 position is cytidine, the binding pocket appears to be flexible enough to accommodate an adenine at this position. During the processive scanning of ssDNA, the binding pocket likely allows binding of all nucleotides before encountering the preferred hotspot sequence for catalysis. The conformation we captured therefore possibly reflects an intermediate scanning state, but not the catalytic state.

In our structure, the adenosine is stabilized by a network of hydrogen bonds and stacking interactions in the −1 pocket (Fig 2A and B). The backbone of residues P210 and I314 hydrogen bond with the amine group of the adenine, while Y315 hydrogen bonds with the phosphate group of the DNA backbone. The adenine base is further stabilized in the pocket through stacking interactions with W211, W285, and Y315 (Fig 2B). The overall architecture of the A3G_CTD_ DNA binding pocket is conserved when compared to other structures of A3s bound to DNA. Our structure aligns well with the structures of A3A bound to thymidine at the −1 position (PDBIDs 5SWW, 5KEG) (31, 32) with an overall root mean square deviation (RMSD) of 0.8 Å and 0.7 Å, respectively. Comparison between the A3A and A3G_CTD_ DNA bound structures reveals that highly conserved residues within the A3 family (A3G residues Y315 and W285) adopt similar orientations in the −1 pocket to bind to a nucleotide (Fig 2C). This highlights that some interactions between the −1 pocket and DNA are conserved between the proteins in the A3 family, even though different nucleotides are preferred at this position.

**Fig 2.**
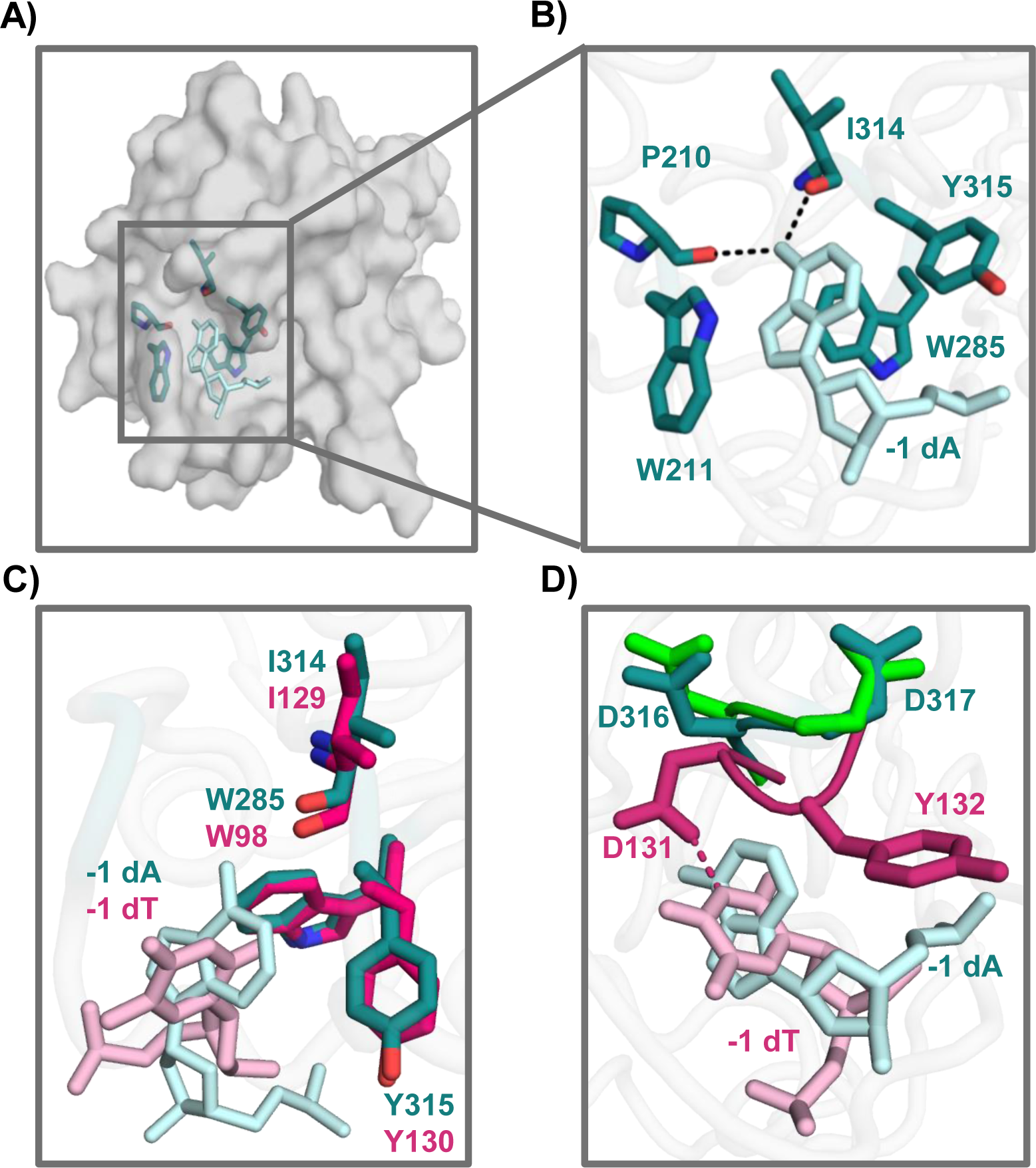
Structural analysis of loop 7 in A3G-ssDNA interaction. A) Overview of the A3G_CTD_, shown in gray surface representation, bound to an adenine in the −1 nucleotid-binding pocket, shown in light blue sticks. Selected A3Gctd residues in the pocket are shown in teal sticks. B) The backbones of I314 and P210 form hydrogen bonds with the adenosine (shown in light blue) in the −1 nucleotide pocket. W211, W285, and Y315 stack with the nucleotide to stabilize it in the pocket. C) Structural alignment of the A3G_CTD_ (teal) to A3A (magenta, PDBID 5SWW) (31), shows that conserved residues W285, I314, and Y315 of the −1 nucleotide-binding pocket are held in similar positions. D) Comparison of the A3G_CTD_, teal, to the A3A-DNA complex, magenta (PDBID 5SWW) (31). In the A3A structure, D131 forms a hydrogen bond with the Watson-Crick edge of the hotspot nucleotide thymidine (light pink). In the A3G_CTD_ structure, D316 and D317 are flipped 180 degrees to avoid clashing with the non-preferred adenine (light blue). The flipped conformation is similar to that of the apo structure of the A3G_CTD_, green (PDBID 3IR2) (23).

Analysis of the adenine-bound A3Gctd structure in comparison to other A3 crystal structures reveals that residues in the −1 pocket change conformations in response to the identity of the nucleotide that A3G encounters. Previous studies have shown that A3G_CTD_ loop 7, specifically residue D317, is important for −1 nucleotide selectivity (29). When comparing our structure of A3G_CTD_ to the A3A-DNA structures (PDBID 5SWW, 5KEG) (31, 32), residues D316 and D317 change their conformations. When an A3A-preferred thymine nucleotide is bound to the −1 pocket, D131 of A3A (corresponding to D136 in A3G) forms a hydrogen bond with the Watson-Crick edge of the thymidine to orient the nucleotide in the pocket (Fig 2D). If A3G_CTD_ D316 remained in a similar orientation as A3A D131, this residue would sterically clash with the adenine base. Instead, the A3Gctd −1 pocket accommodates the adenine base by flipping D316 180 degrees away from the nucleotide (Fig 2D), which opens the −1 pocket to allow for adenine to bind. To compensate for the loss of the hydrogen bond between the base and D316, A3G_CTD_ interacts with the Watson-Crick edge of the adenine through backbone hydrogen bonds (Fig 2B). The adenine bound structure of A3G_CTD_ aligns to the apo structures of A3G_CTD_ (PDBIDs 3IR2 (23),4ROW/4ROV (36), 3IQS/3E1U (21)) with an RMSD of ~0.4 Å. Residues D316 and D317 in the adenine bound structure adopt a similar conformation to that of the apo structure (Fig 2D, green residues). This suggests that A3G samples this conformation in the absence of DNA and while scanning the DNA for its preferred hotspot sequence.

### A3G loop 1 acts as the scanning module for −1 nucleotide recognition

In addition to the conformational changes of loop 7 residues, we also observed structural changes of A3G_CTD_ loop 1. Comparison of our adenine bound structure to the A3G_CTD_ apo structures (PDB 3IR2 (23),4ROW/4ROV (36), 3IQS/3E1U (21)) reveals that loop 1 of A3G_CTD_ swings approximately 3Å towards the adenine (Fig 3A). The movement of loop 1 leads to the repositioning of W211 for stacking interactions with the adenine as well as bringing P210 closer to the base to allow for a hydrogen bond to form with the nucleotide (Fig 3B). Notably, while W211 on loop 1 flips inwards to close the binding pocket upon adenine binding, residues Y315 and W285 remain static (Fig 3B). The conformational changes that loop 1 undergoes in response to binding a non-preferred nucleotide in the −1 pocket suggests that this loop is also a flexible protein module, allowing for nucleotides that differ from the hotspot sequence to enter the nucleotide-binding pocket during scanning.

**Fig 3.**
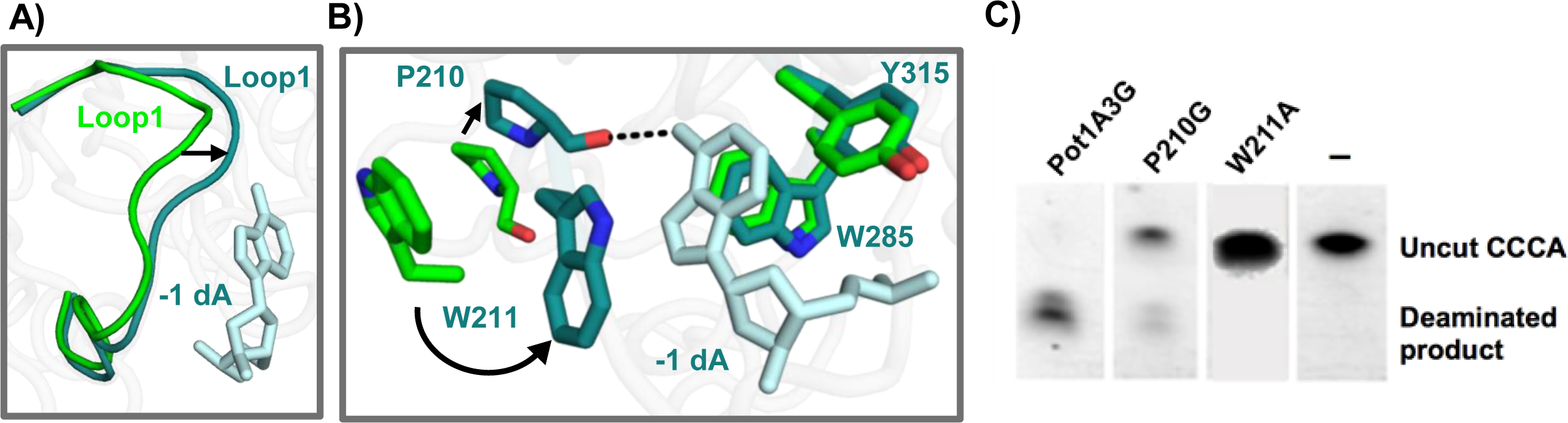
A3G_CTD_ loop 1 is important for substrate recognition. A) Loop 1 in A3G_CTD_-DNA complex (teal, coil representation) moves 3Å compared to apo A3G_CTD_ (PDBID 3IR2, green) (23) to enclose the adenine in the −1 pocket. B) Comparison of the A3G_CTD_ apo crystal structure (green; PDB ID 3IR2) (23) to the A3G_CTD_-DNA structure (teal). Residue W211 flips in to stack with the nucleotide and residue P210 moves toward the nucleotide as compared to the apo structures, while W285 and Y315 remain static. C) Deaminase activity assays on A3Gctd mutants shows that mutating residues on loop 1 can disrupt the deaminase activity of A3G completely in the case of W211A or partially in the case of P210G.

To further confirm the importance of loop 1 in catalysis, we mutated residues P210 and W211 and determined the enzymatic activity of the mutants. We found that mutating W211 to alanine abolished the catalytic activity of the enzyme (Fig 3C). This is consistent with the previous mutagenesis findings that the aromatic residues in the −1 nucleotide-binding pocket are critical for the catalytic and antiviral activity of A3G (20, 21, 23). Furthermore, as proline residues introduce conformational constraints to the protein backbone, we examined the importance of rigidity of loop 1 by mutating P210 to a flexible glycine. This mutation caused a significant reduction in the deaminase activity (Fig 3C). These results suggest that loop1 is important not only for interacting with the nucleotides, but also affects A3G catalysis.

### Mutating the −1 nucleotide-binding site of A3G affects hotspot preference at the +1 position

The recent A3A-DNA structures (31, 32) show that the ssDNA substrate is bound to the protein in a U-shaped conformation, with the +1 and the −1 nucleotide-binding pockets positioned adjacent to each other (Fig 4A). The close clustering of the nucleotide-binding pockets may allow the binding sites to influence each other. To examine if these sites are interconnected, we tested whether mutating −1 nucleotide-binding residues in loop 1 would affect the A3G hotspot preference at the +1 position. Some of the amino acid residues forming the −1 nucleotide-binding pocket (W285, I314, and Y315) are highly conserved among the human A3 family and AID (Fig4A). In contrast, the A3 proteins and AID vary in sequence in loop 1 at residues P210 and W211 in the A3G_CTD_; those with proline at the P210-equivalent positions favor adenine at the +1 site (A3G and A3B) (2), while those with arginine at this position have a higher propensity to favor a thymidine at this position (AID and A3F), although A3F still has a slight preference for an adenine (2, 37) (Fig 4B). This suggests that these −1 nucleotide-binding residues may also affect the selectivity for the +1 nucleotide in the ssDNA substrate. Specifically, among the A3 family homologs, AID shares the greatest degree of sequence conservation with A3G in the residues that interact with the +1 base, with only a single amino-acid variation at the A3G P210 (AID R19) position. This single residue may have substantial influence on the DNA preference of the two enzymes for the +1 position, where A3G favors adenosine while AID favors thymidine (38, 39).

**Fig 4.**
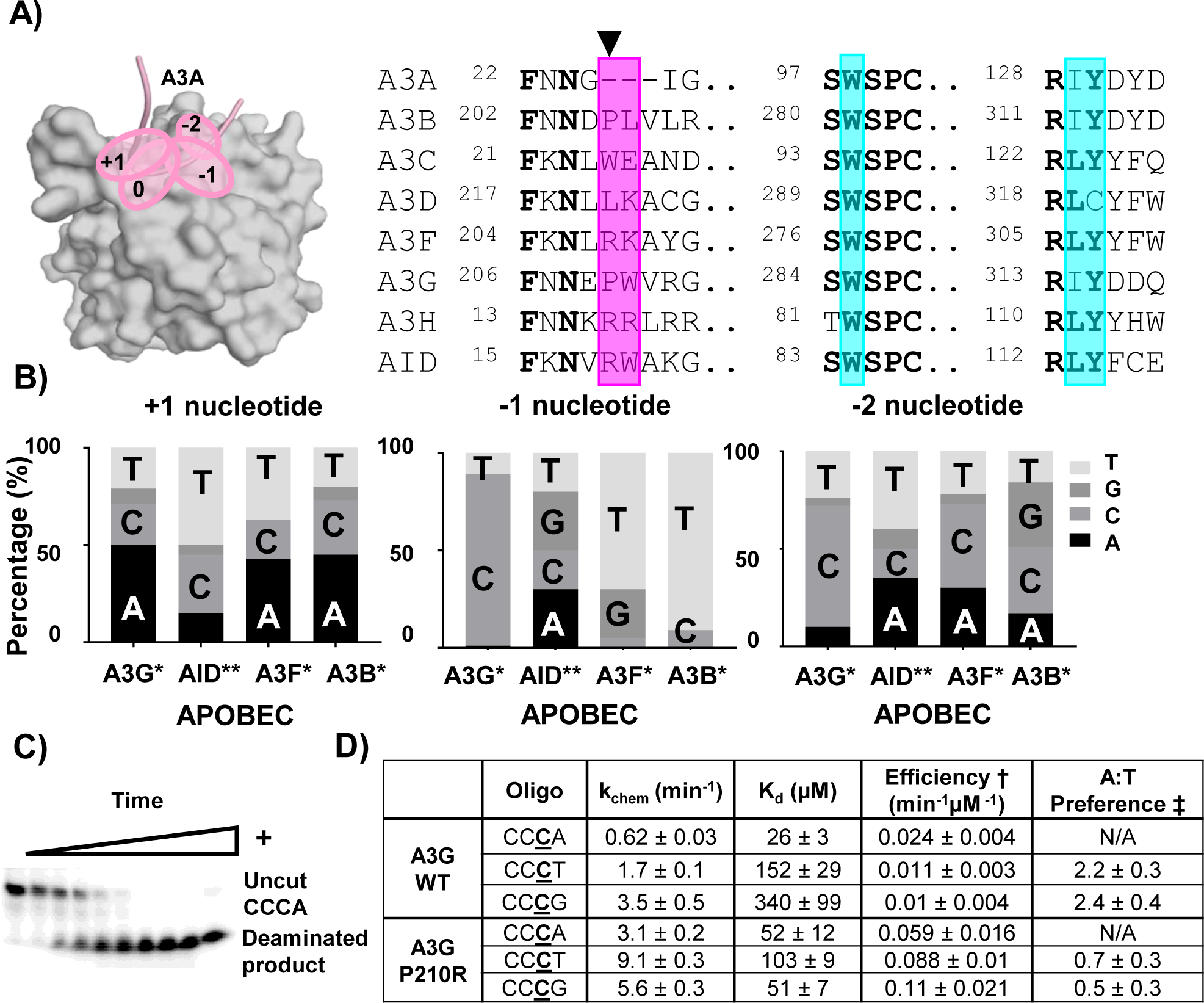
Mutating residues in loop 1 results in change of preference for +1 nucleotide. A) Schematic of the nucleotide binding sites of A3A (PDBID 5SWW) (31) that are spatially close to one another (marked by pink ovals). A3A is in surface presentation and the backbone of the bound ssDNA is shown as a pink coil. Right panel: sequence alignment of the proteins from the A3 superfamily. Conserved residues are in bold. Residues involved in hydrogen bonding with the nucleotide at the −1 position during scanning are shaded in magenta, and the other residues forming the −1 nucleotide pocket are shaded in teal. The arrow marks the A3G P210 corresponding position. B) Frequency (%) of the preference for each nucleotide at the −2, −1, and +1 positions for A3B, A3F, A3G (2) and AID (37). C) ^32^P-labeled oligonucleotides were used in a UDG-dependent cleavage assay to measure cytidine deaminase activity of Pot1A3G_CTD_. D) A3G_CTD_ catalytic parameter measurements and sequence preference as determined by a UDG-dependent cleavage assay (graphs shown in supplementary figure 1). Error values are based on fits to the hyperbolic K_d_ curve, *kobs* = *(kchem*[E])/(K_d_+[E])*. The errors represent standard errors of the parameters. † Efficiency = k_chem_/K_d_, ‡ Preference = Efficiency(CC**<u>C</u>**A)/Efficiency(CC**<u>C</u>**T/G)

We examined the extent to which the identity of the amino acid at the A3 G P210 position dictates the +1 nucleotide preference. To mimic AID, we mutated P210 in A3G to arginine and measured cytidine deamination kinetics and substrate DNA preference using different ssDNA oligonucleotides containing a solitary target cytidine (Fig 4C). We conducted single-turnover kinetics experiments to determine the K_d_ and k_chem_ for WT and P210R for different ssDNA oligonucleotides (Fig 4D and S1 Fig), and found that WT A3G_CTD_ binds DNA with the canonical hotspot with a Kd of 26 μM (Fig 4D). Notably, the P210R mutation decreased the affinity for CC**<u>C</u>**A, while increasing the affinity for both CC**<u>C</u>**T and CC**<u>C</u>**G substrates. The preference for the +1 nucleotide was calculated based on the ratio of efficiency for catalyzing DNA containing adenine compared to thymine or guanine at this position. A3G_CTD_ preference for adenosine at the + 1 position was consistent with previous observations (2, 37). However, the A3G_CTD_ P210R mutant had a three-fold decrease in A:T preference (from 2.2 to 0.7) compared to WT, indicating a switch of preference from adenosine to thymidine at the +1 position. This is consistent with the fact that the AID, which prefers thymidine in the +1 position, has an arginine at the corresponding residue in loop 1. Together, these results show that mutations in the-1 nucleotide-binding pocket perpetuate changes in the +1 nucleotide preference, supporting that the two nucleotide-binding pockets are not independent, but are structurally connected and communicate with each other.

### Mutating loop 1 of A3G has a broad effect on hotspot mutation rates *in vivo*

To confirm that mutating the −1 pocket affects the preference for the other nucleotide positions in the hotspot, we evaluated the full-length A3G P210R mutant on both its antiviral activity and the sequence context of the G-to-A hypermutation in human cell culture. We produced VSV-G pseudotyped viruses in HEK293T cells in the presence of A3G-WT or A3G-P210R and determined infectivity in TZM-bl cells in a single round of replication, as previously described (40) (Fig 5A). In line with previous studies (1, 5, 24, 41), WT and mutant A3G potently blocked replication of HIV-1 produced in their presence compared to viruses generated in their absence (Fig 5A). The presence of viral infectivity factor (Vif) completely averted both WT and P210R mutant A3G antiviral activity. These results demonstrated that the P210R substitution did not affect the overall antiviral activity of A3G or its susceptibility to Vif-mediated degradation.

**Fig 5.**
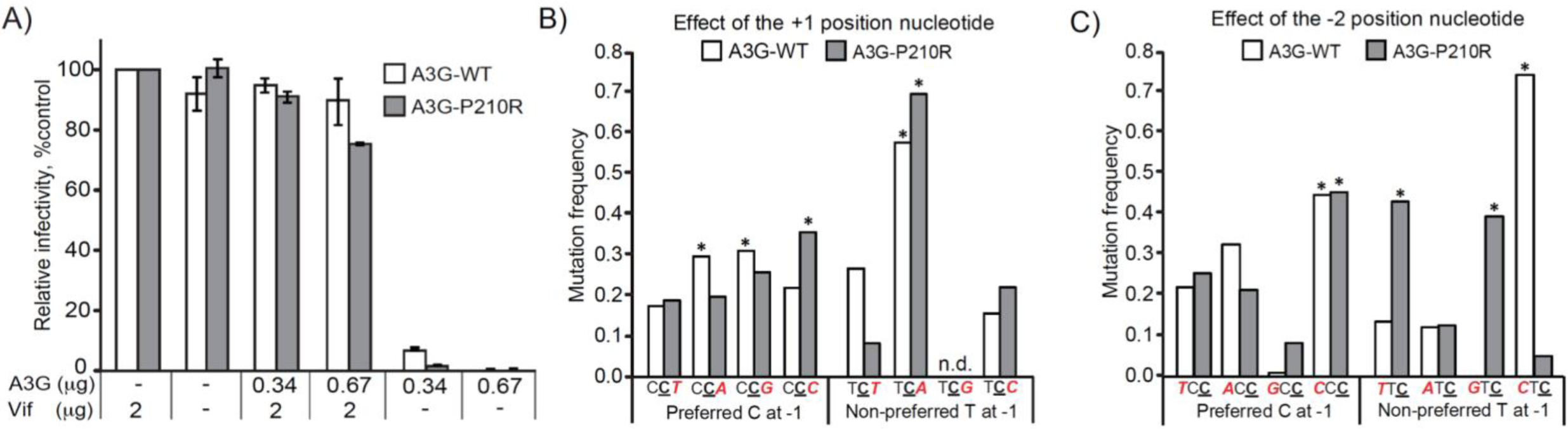
P210 is important for selection of the −2 and +1 nucleotides at the A3G deamination sites. A) Relative single-cycle infectivity of VSV-G-psuedotyped HIV-1 Δ*vif*viruses produced in the presence or absence of A3G-WT or A3G-P210R. Mean of two independent experiments done in triplicates are shown relative to the “no A3G” control (set to 100%); error bars, standard deviation. B and C) Effect of A3G-P210R substitution on the relative mutation frequencies at the +1 position nucleotide (b) and the −2 position nucleotide (c) with the preferred C at the −1 position (5’C**<u>C</u>**) or the non-preferred T at the −1 (5’T**<u>C</u>**) position. B) In the + 1 position nucleotide, mutation frequencies for the preferred sites with a C at −1 position (5’C**<u>C</u>**), A3G-WT prefers C**<u>C</u>**A or C**<u>C</u>**G over C**<u>C</u>**T and C**<u>C</u>**C; A3G-P210R prefers C**<u>C</u>**C over C**<u>C</u>**A, C**<u>C</u>**G, and C**<u>C</u>**T, but has no significant difference in preference between C**<u>C</u>**A and C**<u>C</u>**T. For the non-preferred sites with a T at −1 position (5’T**<u>C</u>**), A3G-WT and A3G-P210R both prefer T**<u>C</u>**A over T**<u>C</u>**T, TCG, or T**<u>C</u>**C. C) In the −2 position nucleotide, mutation frequencies for the preferred sites with a C at −1 position (5’C**<u>C</u>**), both A3G-WT and A3G-P210R prefer *CCC* over TC**<u>C</u>**, AC**<u>C</u>**, or GC**<u>C</u>**. For the non-preferred sites with a T at the −1 position (5’T**<u>C</u>**), A3G-WT prefers CT**<u>C</u>** over TT**<u>C</u>**, AT**<u>C</u>**, or GT**<u>C</u>**, whereas A3G-P210R prefers TT**<u>C</u>** or GT**<u>C</u>** over AT**<u>C</u>** and CT**<u>C</u>**. The significantly preferred nucleotide in the −2 or +1 positions are indicated (\**P*< 0.001). The number of sites, number of mutations, and relative mutation frequencies are shown in Supplemental Table 1.

Despite a comparable overall antiviral activity of WT A3G and the P210R mutant, our biochemical data suggest that each A3G construct exhibits different nucleotide preference at +1 position, even though P210 is interacting with the −1 nucleotide (Fig 4D). We therefore infected CEM-SS T cells with virus produced in the presence of A3G-WT or A3G-P210R, and analyzed G-to-A hypermutation patterns in sequences of proviral DNAs (Fig 5B). The overall 5’GG-to-AG hypermutation frequency was similar for both WT and P210R A3G, indicating that the P210R mutation did not significantly alter the deamination activity of the enzyme; however, substrate selectivity was altered for the +1 position. WT A3G preferred adenosine at the +1 position (5’C**<u>C</u>**A:5’C**<u>C</u>**T ratio = 0.28/0.17; P < 0.0001), whereas the P210R mutant had a substantial decrease in its preference for an adenosine (5’C**<u>C</u>**A:5’C**<u>C</u>**T ratio = 0.11/0.10; P > 0.05) (Fig 5B, S1 Table). This change in 5’C**<u>C</u>**A:5’C**<u>C</u>**T preference is consistent with, but not as drastic as observed in our biochemical studies using the A3G_CTD_ (Fig 4D), likely because the A3G_NTD_ also interacts with the DNA (42) and thus, contributes to substrate specificity. Nonetheless, the substantial change in 5’C**<u>C</u>**A:5’C**<u>C</u>**T ratio in the cell-based assay using full-length A3G variants corroborates the notion that the nucleotide binding pockets are tightly entwined.

Mutating P210 also affects the nucleotide preference at the −2 position in the context of the less preferred thymidine at the −1 position (5’T**<u>C</u>**), but not in the context of the preferred cytidine at the −1 position (5’C**<u>C</u>**). Both WT A3G and P210R A3G preferred cytidine compared to thymidine at the −2 position when the −1 position was a cytidine (WT 5’CC**<u>C</u>**:5’TC**<u>C</u>** ratio = 0. 37/0.18; P < 0.0001 and P210R 5’CC**<u>C</u>**;:5’TC**<u>C</u>** ratio = 0.21/0.12; P < 0.0001) (Fig 5C, S1 Table). When the −1 nucleotide was the less preferred thymidine, WT A3G maintained a preference for cytidine at the −2 position (5’CT**<u>C</u>**:5’TT**<u>C</u>** ratio = 0.07/0.01; P < 0.0001) (Fig 5C, S1 Table). In contrast, P210R had a substantially increased preference for thymidine and guanidine compared to cytidine at the −2 position (5’TT**<u>C</u>**:5’CT**<u>C</u>** ratio = 0.05/0.01; P < 0.0001 and 5’GT**<u>C</u>**:5’CT**<u>C</u>** ratio = 0.05/0.01; P < 0.0001) (Fig 5C, S1 Table). In summary, changes to the −1 nucleotide pocket, specifically the P210 residue, affect the nucleotide preference at the +1 position when the −1 nucleotide is the preferred cytidine (5’C**<u>C</u>**) and at the −2 position when the - 1 nucleotide is the non-preferred thymidine (5’T**<u>C</u>**). Therefore, these results lend support to our biochemical and structural studies with Pot1A3G_CTD_, demonstrating that changes to the −1 nucleotide pocket, specifically the P210 residue, affect the entirety of the hotspot preference of A3G.

## Discussion

A3G is one of the most potent restriction factors of HIV-1, yet its mechanism of substrate selection is still poorly understood. A3G prefers to deaminate cytidines in the hotspot sequence 5’-CC**<u>C</u>**A (where the deaminated cytidine is underlined) (2, 26). Despite this hotspot preference, deamination can still occur to a lesser extent when other nucleotides are at the flanking positions. For example, A3G is capable of cytidine deamination with any nucleotide at the +1 position, albeit at different frequencies as shown in Figs 4 and 5B/C. In addition, it has been shown that the cytidine at the −1 position (the 5’ side of the deaminated cytidine) can also be deaminated (43). In this study, we used the novel fusion of Pot1 to the A3G_CTD_ to capture the low affinity A3GcTD-ssDNA interaction, identifying a non-preferred adenosine in the −1 nucleotide-binding pocket of A3G_CTD_. This is the first structure of an A3 bound to a non-preferred hotspot substrate. Since A3G is a highly processive enzyme (35), it frequently encounters purines in the −1 nucleotide binding pocket while scanning the DNA for its hotspot. It was unknown how A3G allows binding but discriminates against deaminating such substrates.

Our comparative structural analysis, biochemistry, and virology studies provide insight into how encountering a purine in the A3G −1 pocket would not result in deamination of a cytidine at the 0 position. We show that unlike the A3A-DNA interactions with the preferred substrate, residue D316 does not flip in to interact with the Watson-Crick edge of the base (Fig 2D) and there is no selectivity toward the nucleotide. A3G instead uses the backbone of P210 in loop 1 to interact with the −1 nucleotide along the Hoogsteen edge of the non-preferred adenine, which causes a structural change in the A3G −1 nucleotide-binding pocket (Fig 3A). This structural change further perturbs the conformation of other nucleotide-binding sites; as recent structures (31, 32) and our biochemical and cell-based data show, these sites are clustered and inter-connected. In fact, mutating residues ^210^PW^211^ causes deformations in the −1 nucleotide pocket that perpetuate preference changes throughout the entire hotspot sequence (Figs 4 and 5). Thus, these perturbations from binding non-preferred nucleotides in the −1 nucleotide pocket may shift the residues in the catalytic pocket making the local environment non-ideal for the deamination reaction.

Our analysis provides an understanding toward the A3G scanning state that allows it to find its preferred deamination site in DNA. During the process of DNA scanning, when A3G encounters a non-preferred sequence, the rearrangements in the binding pockets necessary to accommodate the nucleotide result in a conformation that is not suitable for deamination at the catalytic site (Fig 6B). Only when the enzyme encounters the hotspot sequence, the collaborative interactions of the preferred nucleotides and their binding pockets on A3G result in a catalytically productive conformation for deamination (Fig 6A).

**Fig 6:**
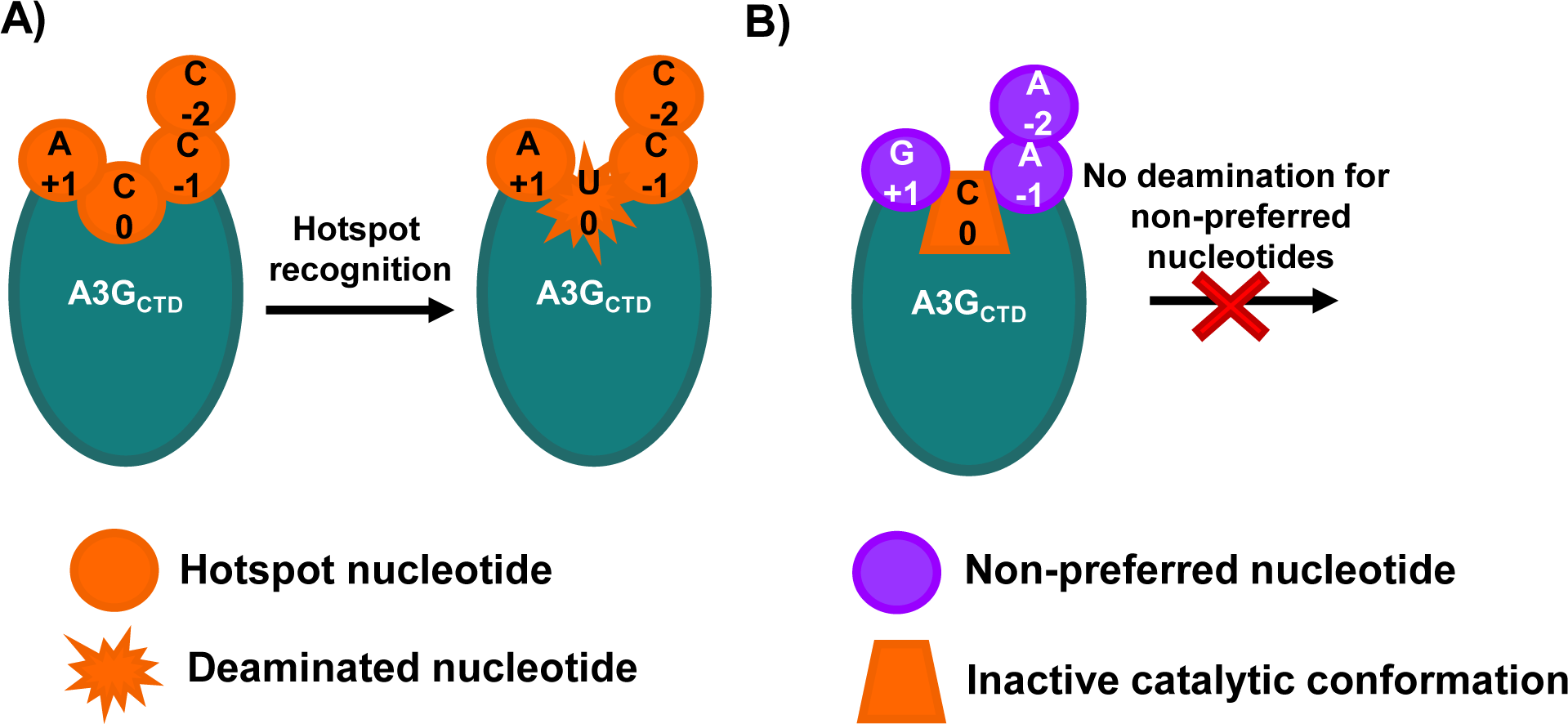
Schematic for DNA selection and nucleotide pocket communication. A) When A3G (represented by a teal oval) encounters a hotspot (preferred nucleotides represented by orange circles), the A3G is active and the cytidine in the 0 position is deaminated, resulting in a uridine at the 0 position (orange star). B) When A3G (teal oval) encounters non-preferred nucleotides flanking a cytidine (purple circles), it adapts an unfavorable conformation (orange trapezoid) at the catalytic site and no deamination occurs.

## Materials and Methods

### Protein expression and purification

His_6_-Staph Nuclease (SN) with SARS-CoV Mpro cleavage site (44, 45) was inserted into the NcoI - BamHI site of pRSF-Duet-1 plasmid (Novagen), followed by the insertion of Pot1-A3G195-384-2K3A fusion gene into the EcoRI – XhoI site (22, 34, 46). The wild-type A3G_CTD_ was constructed from A3G191-384-2K3A gene inserted into NcoI - XhoI site of the expression vector pMAT9s, containing an N-terminal His_6_-tag followed by maltose binding protein (MBP) and a SARS-CoV Mpro cleavage site (44). All A3G mutants, including P210R and W211A were generated using this construct as a template. The vectors encoding A3G_CTD_ mutants were made by QuickChange mutagenesis. All constructs were transformed and expressed in BL21 (DE3) *Escherichia coli*cells grown in TB media to an OD600 of 0.6 and induced with 0.5 mM isopropyl β-D-1-thiogalactopyranoside (IPTG) overnight at 16°C. The cells were then harvested by centrifugation (5000 rpm, 10 min, 4°C) and resuspended in lysis buffer [50 mM Tris (pH 7.5), 500mM NaCl, and 0.1mM Tris(2-carboxyethyl)phosphine (TCEP)]. Resuspension was followed by lysis with a microfluidizer. The lysate was centrifuged [13,000 rpm, 40 min, 4°C] and proteins were purified by nickel affinity column (Qiagen) on FPLC (GE Healthcare). SN or MBP tag was removed by digestion with SARS-CoV Mpro protease overnight at 4°C. The target protein was separated using a HiTrapQ anion exchange column (GE Heathcare) in 20 mM Tris (pH 8.0) using a 0 - 1000 mM NaCl (with 0.1mM TECP) gradient elution, followed by Superdex-200 gel-filtration column (GE Heathcare) in corresponding buffer [50 mM Tris (pH 7.0) (for Pot1-A3G_CTD_) or 50 mM NaH_2_PO_4_ (pH 8.0) (for all A3G_CTD_ and mutants), 100 mM NaCl and 0.1 mM TCEP]. The protein purity was examined by SDS-PAGE. Pot1-A3G_CTD_ was mixed with a DNA oligonucleotide [30nt substrate: AGA AGA CC**<u>C</u>** AAA GAA GAG GAA GCA GGT TAC] at 1:1 molar ratio, and further purified using a Superdex-75 size-exclusion column. The protein/DNA complex was then concentrated to 3 mg/mL for crystallization.

### Crystallization and data collection

Pot1-A3G_CTD_/DNA complex crystals were grown at 20°C using the microbatch-under-oil method by mixing equal amounts of sample [in buffer of 50 mM Tris (pH 7.0), 100 mM NaCl and 0.1 mM TCEP] and crystallization buffer [100 mM HEPES pH7, 200 mM LiCl, and 20% (w/v) Polyethylene glycol (PEG) 6000]. Crystals were cryo-protected by the crystallization buffer with 30% (v/v) PEG 400 and frozen in liquid nitrogen. Diffraction data were collected at the National Synchrotron Light Source beamline X29A to the resolution of 2.9 Aࡪ. Data were processed using HKL2000 (47). Analysis of the data showed that the crystal has close to perfect twinning. The data statistics are summarized in Table 1. Coordinates and structural factors have been deposited in the Protein Data Bank under the accession code 6BWY.

### Structure determination and refinement

There were four Pot1-A3G_CTD_ molecules in the asymmetric unit of the crystal. The structure was solved by molecular replacement using PHASER (48) with the A3Gctd structure (PDBID 3IR2 (23)) and the Pot1 structure (PDBID 1QZH (34)) as search models. Clear electron density of Pot1-A3G_CTD_, including that for the Pot1 cognate DNA, was evident in the electron density map. Additional electron density was observed for the adenine at the 5’ side of the Pot1 cognate DNA sequence. Furthermore, weak electron density 5’ to the density was also observed but the quality was not sufficient for model building. Rigid body and iterative rounds of restrained refinement (including amplitude-based twin refinement) were carried out using Refmac5 (49), followed by rebuilding the model to the 2Fo-Fc and the Fo-Fc maps using Coot (50). Non-crystallographic symmetry restraints were applied in the refinement cycles. The final model has an R_work_/R_free_ of 23.3%/28.9%. The refinement statistics are summarized in Table 1. The structure was analyzed and illustrated with Coot and PyMOL (51).

### Deaminase assay using fluorescent-tagged ssDNA substrates

The substrate 30-mer oligos containing CC**<u>C</u>**A, CC**<u>C</u>**T, and CC**<u>C</u>**G (IDT, Coralville, IA) were labeled with 6-carboxyfluorescein (6-FAM) at the 5’ terminus. 2 nM 6-FAM-labeled oligos were incubated at 37°C for different time lengths, with 30μg A3G protein samples and 5 units of Uracil-DNA Glycosylase (UDG) (New England BioLabs, Ipswich, MA). The abasic sites were then hydrolyzed by a 30-minute incubation with 0.25 M NaOH, which was followed by the addition of 20 μL of 1M Tris-HCl, pH 8.0. The reaction products were separated on a TBE-Urea PAGE gel (Life Technologies, Carlsbad, California). Gel bands were imaged with a CCD imager.

### Radiolabeling of primers for *in vitro* kinetics

The substrate 30-mer oligos CC**<u>C</u>**A, CC**<u>C</u>**T, and CC**<u>C</u>**G (IDT, Coralville, IA) were radiolabeled at the 5’ terminus with [*γ*-^32^P] ATP (Perkin Elmer, Waltham, MA) using T4 polynucleotide kinase (New England Biolabs, Ipswich, MA), as described previously (35, 52). Radiolabeled oligos were desalted using a Bio-Spin 6 column (Bio-Rad Laboratories, Hercules, CA).

### A3G *in vitro* single-turnover kinetics

WT or P210R A3G_CTD_ was buffer exchanged into Reaction Buffer (20 mM Tris pH 8.0, 1 mM DTT). Enzyme at varying concentrations (1 μM – 50 μM) was incubated at 37°C for 5 minutes and reactions were induced with 40 nM of ^32^P-labeled DNA oligomer. At given time points, 8 μL of the enzyme-oligo mixture was removed from the mix and quenched by the addition of 12 μL of quench buffer (50 mM EDTA, pH 8.0, final concentration) preheated to 95°C. After a 5-minute incubation at 95°C, the quenched mixture was incubated at 37°C for 5 minutes. Five units of UDG (New England BioLabs, Ipswich, MA) were incubated with the A3G-quenched mixture for two hours to cleave free uracil from any uracil-containing oligomers formed by A3G catalysis. The abasic sites were then hydrolyzed by a 30-minute incubation with 0.25 M NaOH, which was followed by the addition of 20 μL of denaturing PAGE dye.

The reaction products were separated on a 20% denaturing PAGE gel. Gel band intensities were measured by Bio-Rad Phosphorimager (Bio-Rad Laboratories) and analyzed by Quantity One software (Bio-Rad Laboratories). The ratio of the intensities of cleaved to uncleaved oligomer at each time point were plotted using Kaleidagraph (version 4.03, Synergy Software) and the rate at a given concentration of enzyme was fit to a single exponential curve, *Percent converted = A(1-e^−kobs*time^)*, where A is maximum conversion (~100%) and k_obs_ is the single-turnover rate. The resultant rates at varying concentrations of enzyme were plotted using Kaleidagraph and fit to a hyperbolic K_d_ curve, *k_obs_* = *(k_chem_*[E])/(K_d_+[E])*, where k_chem_ is the rate of chemistry and K_d_ is the dissociation constant. Errors given are standard errors of parameters.

### Cell culture, Plasmids, transfections, and virus production

HEK293T, TZM-bl, and CEM-SS cell lines were obtained from the American Type Culture Collection. HEK293T and TZM-bl cell lines were maintained in Dulbecco’s modified Eagle’s medium and CEM-SS cell line was maintained in RPMI 1640 medium (Corning Cellgro). Both media were supplemented to contain 10% fetal calf serum (Hyclone), 100 IU/ml penicillin and 100 μg/ml streptomycin (GIBCO).

A3G-P210R mutant was generated by site-directed mutagenesis (QuickChange Lightening site-directed mutagenesis kit, Agilent Technologies) using the following primers: P210R_sense:

5’ATTCACTTTCAACTTTAACAATGAACGGTGGGTCAGAGGAC3 ’ P210R_antisense:

5’ GTCCTCTGACCCACCGTTCATTGTTAAAGTTGAAAGTGAAT3 ’.

All viruses were prepared using a previously described HIV-1 vector pHDV-eGFP pseudotyped by co-transfecting with phCMV-G plasmid, which expresses vesicular stomatitis virus glycoprotein (VSV-G) (41, 53–58). Briefly, we co-transfected pHDV-eGFP (1.0 μg), pHCMV-G (0.25 μg), and either 0.34 μg or 0.67 μg of pFlag-WT-A3G or pFlag-P210R-A3G expression plasmids in the presence or absence of pcDNA-hVif using polyethylenimine (PEI) as previously described (41, 53–55). Virus-containing supernatant was clarified by filtering through a 0.45-μm filter and kept at −80°C until use.

### Virus infectivity and hypermutation analysis

Virus p24 CA amounts were determined using enzyme-linked immunosorbent assay (XpressBio). TZM-bl indicator cells (59) were infected using equivalent p24 CA amounts of viruses, and infectivity was determined by measuring luciferase enzyme activity using Britelite luciferase solution (PerkinElmer) and a LUMIstar Galaxy luminometer (PerkinElmer).

For the hypermutation pattern analysis, CEM-SS cells (300,000 cells/well in 24-well plate) were infected with HIV-1 Δ*vif*-virus produced in the presence of 0.67 μg A3G. Cell pellets were collected 72 h post-infection and total DNA was extracted using QIAamp DNA blood minikit (Qiagen Inc). The reverse transcriptase (RT) coding region of HIV was PCR amplified using primers (HIV-10 FW: GGACAGCTGGACTGTCAATGACATAC and HIV-11 rev: GTTCATTTCCTCCAATTCCTTTGTGTG) and cloned into the pCR2.1-TOPO backbone vector using TOPO TA cloning kit (Invitrogen). To avoid bias in our analysis, we selected an 893-nt fragment that has a comparable number of 5’-C**<u>C</u>**A (19 editing sites) and 5’-C**<u>C</u>**T (20 editing sites) trinucleotide hotspots; the fragment also has 61 preferred 5’C**<u>C</u>** editing sites and 76 less preferred 5’T**<u>C</u>** editing sites. Individual clones were purified, sequenced, and analyzed for evidence of A3G-mediated G-to-A hypermutation using the HYPERMUT software (www.hiv.lanl.gov).

### Statistical analysis

All comparisons were made based on the WT-A3G for each mutant using Fisher’s Exact test. Probability values (*P*) < 0.05 were considered significant.

## Acknowledgements

We thank Xiaofang Yu and David Schatz for insightful discussions. We also thank the staff at the Advanced Photon Source beamlines 24ID-C and E, and National Synchrotron Light Source beamline X-25.

## Supporting information

**S1 Fig: Kinetics of the Pot1A3G_CTD_ deamination reaction**

A) Representative kinetic curve to determine the rate of deamination of WT A3G_CTD_. Shown here is the result for 5μM A3G_CTD_.

B) The kinetics plot result for the WT A3G_CTD_ reaction on CC**<u>C</u>**A substrate over a range of A3G concentrations. All other data were analyzed and summarized in Fig4D.

**S1 Table**. Total number of G-to-A mutations was determined for 93 proviruses produced in the presence of A3G-WT and 53 proviruses produced in the presence of A3G-P210R. Hypermutation is defined as ≥ 2 G-to-A mutations per clone (the no A3G control, on average, had <1 G-to-A mutations per clone). Each clone contained 61 5’C**<u>C</u>** and 76 5’T**<u>C</u>** target sites. Mutations/site = total mutations/[sites/clone χ no. of clones]. For all conditions, the mutation frequencies for each nucleotide are shown relative to the total mutations/site as determined by the +1 or −2 position nucleotides. Relative preference of nucleotides at the +1 or −2 position in both the 5’-C**<u>C</u>** and 5’-T**<u>C</u>** edited sites are plotted for virions produced in the presence of A3G-WT or A3G-P210R.

